# The *lhfpl5* ohnologs *lhfpl5a* and *lhfpl5b* are required for mechanotransduction in distinct populations of sensory hair cells in zebrafish

**DOI:** 10.1101/793042

**Authors:** Timothy Erickson, Itallia V. Pacentine, Alexandra Venuto, Rachel Clemens, Teresa Nicolson

## Abstract

Hair cells sense and transmit auditory, vestibular, and hydrodynamic information by converting mechanical stimuli into electrical signals. This process of mechano-electrical transduction (MET) requires a mechanically-gated channel localized in the apical stereocilia of hair cells. In mice, lipoma HMGIC fusion partner-like 5 (LHFPL5) acts as an auxiliary subunit of the MET channel whose primary role is to correctly localize PCDH15 and TMC1 to the mechanotransduction complex. Zebrafish have two *lhfpl5* genes (*lhfpl5a* and *lhfpl5b*), but their individual contributions to MET channel assembly and function have not been analyzed.

Here we show that the zebrafish *lhfpl5* genes are expressed in discrete populations of hair cells: *lhfpl5a* expression is restricted to auditory and vestibular hair cells in the inner ear, while *lhfpl5b* expression is specific to hair cells of the lateral line organ. Consequently, *lhfpl5a* mutants exhibit defects in auditory and vestibular function, while disruption of *lhfpl5b* affects hair cells only in the lateral line neuromasts. In contrast to previous reports in mice, localization of Tmc1 does not depend upon Lhfpl5 function in either the inner ear or lateral line organ. In both *lhfpl5a* and *lhfpl5b* mutants, GFP-tagged Tmc1 and Tmc2b proteins still localize to the stereocilia of hair cells. Using a stably integrated GFP-Lhfpl5a transgene, we show that the tip link cadherins Pcdh15a and Cdh23, along with the Myo7aa motor protein, are required for correct Lhfpl5a localization at the tips of stereocilia. Our work corroborates the evolutionarily conserved co-dependence between Lhfpl5 and Pcdh15, but also reveals novel requirements for Cdh23 and Myo7aa to correctly localize Lhfpl5a. In addition, our data suggest that targeting of Tmc1 and Tmc2b proteins to stereocilia in zebrafish hair cells occurs independently of Lhfpl5 proteins.

## 2 Introduction

The mechano-electrical transduction (MET) complex of sensory hair cells is an assembly of proteins and lipids that facilitate the conversion of auditory, vestibular and hydrodynamic stimuli into electrical signals. Our current understanding is that the proteins of the MET complex consist of the tip link proteins cadherin 23 (CDH23) and protocadherin 15 (PCDH15) at the upper and lower ends of the tip link respectively (Kazmierczak et al., 2007), the pore-forming subunits transmembrane channel-like proteins TMC1 and TMC2 (Kawashima et al., 2011; Pan et al., 2013, 2018), and the accessory subunits transmembrane inner ear (TMIE) and lipoma HMGIC fusion partner-like 5 (LHFPL5) (Cunningham and Müller, 2019; Xiong et al., 2012; Zhao et al., 2014). How these proteins are correctly localized to the sensory hair bundle and assemble into a functional complex is a fundamental question for understanding the molecular basis of how mechanotransduction occurs.

Recent studies have revealed extensive biochemical interactions between the MET complex proteins that are required for the function and / or stable integration of each component in the complex. In particular, LHFPL5 is a central player in MET complex formation, stability, and function. LHFPL5 (a.k.a. tetraspan membrane protein of the hair cell stereocilia / TMHS) is a four transmembrane domain protein from the superfamily of tetraspan junctional complex proteins. This superfamily includes junctional proteins like claudins and connexins, as well as ion channel auxiliary subunits such as transmembrane α-amino-3-hydroxy-5-methyl-4-isoxazole propionic acid receptor (AMPAR) regulatory proteins (TARPs) and the gamma subunits of voltage gated calcium channels. Pathogenic mutations in LHFPL5 are a cause of non-syndromic sensorineural hearing loss in humans (DFNB67), mice and zebrafish (Longo-Guess et al., 2005; Nicolson et al., 1998; Obholzer et al., 2012; Shabbir et al., 2006). LHFPL5 localizes to the tips of stereocilia (Li et al., 2019; Mahendrasingam et al., 2017; Xiong et al., 2012) where it directly interacts with other MET complex components. Co-immunoprecipitation experiments in heterologous cells suggests that LHFPL5 can directly interact with PCDH15 and TMIE (Xiong et al., 2012; Zhao et al., 2014). A structure of the PCDH15 - LHFPL5 complex has also been reported (Ge et al., 2018).

A precise role for LHFPL5 has yet to be defined. One current hypothesis is that LHFPL5 acts as an auxiliary subunit of the MET channel to stabilize the complex, similar to TARPs and the gamma subunits of voltage-gated calcium channels. In mouse cochlear hair cells, PCDH15 and LHFPL5 require each other for stable localization at the tips of stereocilia (Mahendrasingam et al., 2017; Xiong et al., 2012), consistent with their well-defined biochemical interaction. The partial loss of PCDH15 from the stereocilia explains the observed reduction in the number of tip links and the dysmorphic hair bundles in *Lhfpl5^-/-^* hair cells. Interestingly, although experiments have failed to demonstrate a biochemical interaction between LHFPL5 and the TMCs, LHFPL5 is required for the localization of TMC1, but not TMC2, in the hair bundle of mouse cochlear hair cells (Beurg et al., 2015). Consistent with this finding, the researchers identified a residual TMC2-dependent MET current in *Lhfpl5* mutant mice. The basis for the selective loss of TMC1 is not known.

In zebrafish, Lhfpl5a plays a similar role in mechanotransduction as its mammalian counterparts. A mutation in zebrafish *lhfpl5a* was reported in a forward genetic screen for genes required for hearing and balance (Nicolson et al., 1998; Obholzer et al., 2012). The loss of Lhfpl5a disrupts the targeting of Pcdh15a to stereocilia and results in splayed hair bundles (Maeda et al., 2017). However, Lhfpl5a is not required to correctly localize Tmie and vice versa (Pacentine and Nicolson, 2019), in spite of their biochemical interaction in cultured cells (Xiong et al., 2012). Lhfpl5 still localizes to the tips of stereocilia in *Tmc1*/*Tmc2* double mutants (Beurg et al., 2015), as well as *transmembrane O-methyltransferase* (*tomt*) mouse and zebrafish mutants, which fail to traffic TMCs to the hair bundle (Cunningham et al., 2017; Erickson et al., 2017). This phenotype suggests that Lhfpl5 localization does not require the TMCs. However, a number of questions regarding the functions of Lhfpl5 remain unanswered: 1) In zebrafish, what are the molecular requirements for targeting Lhfpl5a to the hair bundle? 2) Is the Lhfpl5-dependent targeting of Tmc1 to the hair bundle an evolutionarily-conserved aspect of their interaction? 3) *lhfpl5a* mutants have defects in auditory and vestibular behaviors, yet sensory hair cells of the lateral line are unaffected (Nicolson et al., 1998). What is the genetic basis for the persistence of lateral line function in *lhfpl5a* mutants?

In this work, we report that teleost fish have two *lhfpl5* genes, *lhfpl5a* and *lhfpl5b*. The zebrafish *lhfpl5* ohnologs are expressed in distinct populations of larval sensory hair cells: *lhfpl5a* in the auditory and vestibular system; *lhfpl5b* in the hair cells of the lateral line organ. Their divergent expression patterns explain why lateral line hair cells are still mechanically-sensitive in *lhfpl5a* mutants. CRISPR-Cas 9 knockout of *lhfpl5b* alone silences the lateral line organ but has no effect on otic hair cell function. We also show that neither Lhfpl5a nor Lhfpl5b are required for Tmc localization in stereocilia. Additionally, we use a GFP-tagged Lhfpl5a to demonstrate that Myo7aa and the tip link proteins Cdh23 and Pcdh15a are required for proper Lhfpl5a localization in otic hair cell bundles. This study reveals the subfunctionalization of the zebrafish *lhfpl5* ohnologs through the divergence in their expression patterns. Furthermore, our work complements previous results from murine cochlear hair cells by highlighting a conserved association between Lhfpl5 and Pcdh15, but also demonstrating novel requirements for Cdh23 and Myo7aa in localizing Lhfpl5a. Lastly, our work indicates that Lhfpl5-dependent localization of Tmc1 is not a universal feature of sensory hair cells and that Lhfpl5 proteins are required for mechanotransduction independently of a role in localizing the Tmc channel subunits to stereocilia.

## 3 Materials and Methods

### 3.1 Ethics statement

Animal research complied with guidelines stipulated by the Institutional Animal Care and Use Committees at Oregon Health and Science University (Portland, OR, USA) and East Carolina University (Greenville, NC, USA). Zebrafish (*Danio rerio*) were maintained and bred using standard procedures (Westerfield, 2000).

### 3.2 Mutant and transgenic fish lines

The following zebrafish mutant alleles were used for this study: *cdh23^nl9^*, *cdh23^tj264^*, *lhfpl5a^tm290d^*, *lhfpl5b^vo35^*, *myo7aa^ty220^*, *pcdh15a^psi7^*, *pcdh15a^th263b^* (Erickson et al., 2017; Ernest et al., 2000; Maeda et al., 2017; Nicolson et al., 1998; Obholzer et al., 2012). The transgenic lines used in this study were *Tg(-6myo6b:eGFP-lhfpl5a)vo23Tg*, *Tg(-6myo6b:tmc1-emGFP)vo27Tg*, *Tg(-6myo6b:tmc2b-emGFP)vo28Tg* (Erickson et al., 2017), and *Tg(-6myo6b:eGFP-pA)vo68Tg*. All experiments used larvae at 1–7 dpf, which are of indeterminate sex at this stage.

### 3.3 Genotyping

Standard genomic PCR followed by Sanger sequencing was used to identify *cdh23^tj264^*, *cdh23^nl9^*, *lhfpl5a^tm290d^*, *pcdh15a^psi7^*, and *pcdh15a^th263b^* alleles. The *lhfpl5b^vo35^* mutation disrupts a MluCI restriction site (AATT) and we are able to identify *lhfpl5b^vo35^* hetero- and homozygotes based on the different sizes of MluCI-digested PCR products resolved on a 1.5% agarose gel. The following primers were used for genotyping:

*cdh23^nl9^*: Fwd – CCACAGGAATTCTGGTGTCC, Rvs – GAAAGTGGGCGTCTCATCAT;
*cdh23^tj264^*: Fwd – GGACGTCAGTGTTCATGGTG, Rvs – TTTTCTGACCGTGGCATTAAC;
*lhfpl5a^tm290d^*: Fwd – GGACCATCATCTCCAGCAAAC, Rvs – CACGAAACATATTTTCACTCACCAG;
*lhfpl5b^vo35^*: Fwd – GCGTCATGTGGGCAGTTTTC, Rvs – TAGACACTAGCGGCGTTGC;
*myo7aa^ty220^*: Fwd – TAGGTCCTCTTTAATGCATA, Rvs – GTCTGTCTGTCTGTCTATCTGTCTCGCT;
*pcdh15a^psi7^*: Fwd – TTGGCACCACTATCTTTACCG, Rvs – ACAGAAGGCACCTGGAAAAC;
*pcdh15a^th263b^*: Fwd – AGGGACTAAGCCGAAGGAAG, Rvs – CACTCATCTTCACAGCCATACAG.

### 3.4 Phylogeny

Lhfpl5 protein sequences retrieved from either NCBI or Ensembl (Supplemental Table 1). Phylogenetic analysis was done on www.phylogeny.fr using T-Coffee (multiple sequence alignment), GBlocks (alignment curation), PhyML (maximum likelihood tree construction with 100 replicates to estimate bootstrap values), and TreeDyn (visualization) (Dereeper et al., 2008).

### 3.5 mRNA *in situ* hybridization

*lhfpl5a* and *lhfpl5b* probe templates were amplified from total RNA using the following primers: *lhfpl5a* Fwd: AATATTGGTGCATAGACTCAAGGAGG; Rvs: GACTCCAAAATGACCTTTTAACAAACGC. *lhfpl5b* Fwd: TGAAGATCAGCTACGATATAACCGG; Rvs: ACTGTGATTGGTGTATTTCCAGC. The inserts were cloned into the pCR4 vector for use in probe synthesis. mRNA *in situ* probe synthesis and hybridization was performed essentially as previously described (Erickson et al., 2010; Thisse and Thisse, 2008). Stained specimens were mounted on a depression slide in 1.2 % low-melting point agarose and imaged on a Leica DMLB microscope fitted with a Zeiss AxioCam MRc 5 camera using Zeiss AxioVision acquisition software (Version 4.5).

### 3.6 Immunofluorescent staining, FM dye labeling, and fluorescence microscopy

Anti-Pcdh15a immunostaining and FM dye labeling of inner ear and lateral line hair cells were performed as previously described (Erickson et al., 2017). All fluorescent imaging was done on Zeiss LSM 700 or LSM 800 laser scanning confocal microscopes. Z-stacks were analyzed using ImageJ (Schneider et al., 2012). All related control and experimental images were adjusted equally for brightness and contrast in Adobe Photoshop CC.

### 3.7 CRISPR-Cas9 knockout of *lhfpl5b*

An sgRNA targeting the *lhfpl5b* sequence 5’-CAACCCAATCACCTCGGAAT-3’ was synthesized essentially as described (Gagnon et al., 2014). For microinjection, 1 µg of sgRNA was mixed with 1 ug of Cas9 protein and warmed to 37°C for 5 minutes to promote the formation of the Cas9-sgRNA complex. Approximately 2 nl of this solution was injected into wild type embryos at the single-cell stage. Efficacy of cutting was determined in two ways: 1) Single larvae genotyping - Amplification of the target genomic region by PCR and running the products on 2.5% agarose gels to assay for disruption of a homogenous amplicon. 2) Assaying for disruption of the mechanotransduction channel in lateral line hair cells by FM 1-43 dye uptake. Larvae displaying a non-wild type pattern of FM 1-43 dye uptake were raised to adulthood. Progeny from in-crosses of F0 adults were exposed to FM 1-43 dye to identify founders. By outcrossing founder adults, we established a line of fish carrying a 5 base pair deletion in the *lhfpl5b* coding region that causes a S77FfsX48 mutation (*lhfpl5b^vo35^*). This mutation disrupts the protein in the first extracellular loop and deletes the final three of four transmembrane helices in Lhfpl5b.

### 3.8 Acoustic startle response

Quantification of the larval acoustic startle response was performed using the Zebrabox monitoring system (ViewPoint Life Sciences, Montreal, Canada) as previously described (Maeda et al., 2017) with the following modifications. Each trial included six larvae which were subjected to two or three trials of 6 acoustic stimuli. For each individual larva, the trial with best AEBR performance was used for quantification. Positive responses where spontaneous movement occurred in the second prior to the stimulus were excluded from analysis. Trials where spontaneous movement occurred for more than 6 of the 12 stimuli were also excluded from analysis.

### 3.9 Microphonics

We performed the microphonic measurements as previously described (Pacentine and Nicolson, 2019). In brief, we anesthetized 3 dpf larvae in extracellular solution (140 mM NaCl, 2 mM KCl, 2 mM CaCl_2_, 1 mM MgCl_2_, and 10 mM 4-(2-hydroxyethyl)-1-piperazineethanesulfonic acid (HEPES); pH 7.4) containing 0.02% 3-amino benzoic acid ethylester (MESAB; Western Chemical). We pinned the larvae with two glass fibers straddling the yolk and a third perpendicular fiber to prevent sliding. For pipettes, we used borosilicate glass with filament, O.D.: 1.5 mm, O.D.: 0.86 mm, 10 cm length (Sutter, item # BF150-86-10, fire polished). Using a Sutter Puller (model P-97), we created recording pipettes with a long shank with a resistance of 10-20 MΩ, after which we beveled the edges to a resistance of 4-5 MΩ using a Sutter Beveler with impedance meter (model BV-10C). For the glass probe delivering the piezo stimulus, we pulled a long shank pipette and fire polished to a closed bulb. We shielded the apparatus with tin foil to completely ground the piezo actuator. To maintain consistent delivery of stimulus, we always pressed the probe to the front of the head behind the lower eye, level with the otoliths in the ear of interest. We also maintained a consistent entry point dorsal to the anterior crista and lateral to the posterior crista. During delivery of stimulus, the piezo probe made light contact with the head. We drove the piezo with a High Power Amplifier (piezosystem jena, System ENT/ENV, serial # E18605), and recorded responses in current clamp mode with a patch-clamp amplifier (HEKA, EPC 10 usb double, serial # 550089). Voltage responses were amplified 1000x and filtered between 0.1 - 3000 Hz by the Brownlee Precision Instrumentation Amplifier (Model 440). We used a 200 Hz sine wave stimulus at 10 V and recorded at 20 kHz, collecting 200 traces per experiment. Each stimulus was of 40 ms duration, with 20 ms pre- and post-stimulus periods. The piezo signal was low-pass filtered at 500 Hz using the Low-Pass Bessel Filter 8 Pole (Warner Instruments). In Igor Pro, we averaged each set of 200 traces to generate one wave response per larva. To quantify hair cell activity, we calculated the amplitude from base-to-peak of the first peak. Larvae were genotyped as described above.

### 3.10 Hair cell counts

*lhfpl5b^vo35/+^* fish were crossed with *Tg(myo6b:eGFP-pA)vo68Tg; lhfpl5b^vo35/+^* fish to produce wild type and *lhfpl5b^vo35^* siblings expressing green fluorescent protein in hair cells. Wild type and *lhfpl5b* mutant larvae were sorted by FM 4-64 labeling (n = 11 each). In experiment 1, larvae (n = 5 each genotype) were fixed in 4% paraformaldehyde at 5 dpf. In experiment 2, larvae (n = 6 each genotype) were imaged immediately. *lhfpl5a^tm290d/+^* fish were in-crossed to produce wild type and *lhfpl5a^tm290d^* siblings. Larvae were sorted based on the auditory and vestibular defects associated with the *lhfpl5a^tm290d^* homozygous mutants. Larvae were labelled with FM 1-43 (n = 6 each). For all experiments, specimens were mounted in low melting point agarose and the L1, MI1, and O2 neuromasts were imaged on a Zeiss LSM 800 confocal. Cell counts were performed using the Z-stack data.

### 3.11 Statistics

Statistical analyses were done using the R stats package in RStudio (R Core Team, 2019; RStudio Team, 2018). P-values of less than 0.05 were considered to be statistically significant. Plots were made with ggplot2 (Wickham, 2016).

### 3.12 Data availability

The raw data supporting the conclusions of this manuscript will be made available by the corresponding author, without undue reservation, to any qualified researcher.

## 4 Results

### 4.1 Teleost fish have ohnologous *lhfpl5* genes

Genes that have been duplicated as a result of a whole genome duplication (WGD) event are known as *ohnologs* (Ohno, 1970; Wolfe, 2000). Due to the teleost-specific WGD, it is not uncommon to find ohnologous genes in teleost fish where other vertebrate classes possess a single gene. The *Danio rerio* (zebrafish) genome contains two *lhfpl5* genes: *lhfpl5a* (ENSDARG00000045023) and *lhfpl5b* (ENSDARG00000056458). To determine if *lhfpl5* duplication is general to the teleost lineage, we queried either GenBank or Ensembl databases to collect Lhfpl5 protein sequences for humans, mice, chick, frogs, and representative sequences from 14 phylogenetic orders of ray-finned fish (Actinopterygii), including 13 orders from the Teleostei infraclass and one from the Holostei infraclass (Supplemental Table 1). The latter (Spotted gar, *Lepisosteus oculatus*) is commonly used to infer the consequences of the teleost WGD, since the Holostei and Teleostei infraclasses diverged before the teleost WGD (Braasch et al., 2016). Phylogenetic analysis of the Lhfpl5 protein sequences supports the idea that duplicate *lhfpl5* genes originated from the teleost WGD event (Figure 1A). In all 13 teleost orders surveyed, there are two *lhfpl5* genes whose protein products cluster with either the *lhfpl5a* or *lhfpl5b* ohnolog groups. Based on available genomic data, there is no evidence that spotted gar fish have duplicated *lhfpl5* genes. Nor is there evidence that the Salmonid-specific WGD (Allendorf and Thorgaard, 1984) lead to further expansion of the *lhfpl5* family. These analyses suggest that the ancestral *lhfpl5* gene was duplicated in the teleost WGD and that the *lhfpl5* ohnologs were retained in all teleost species examined here.

**Figure 1.**
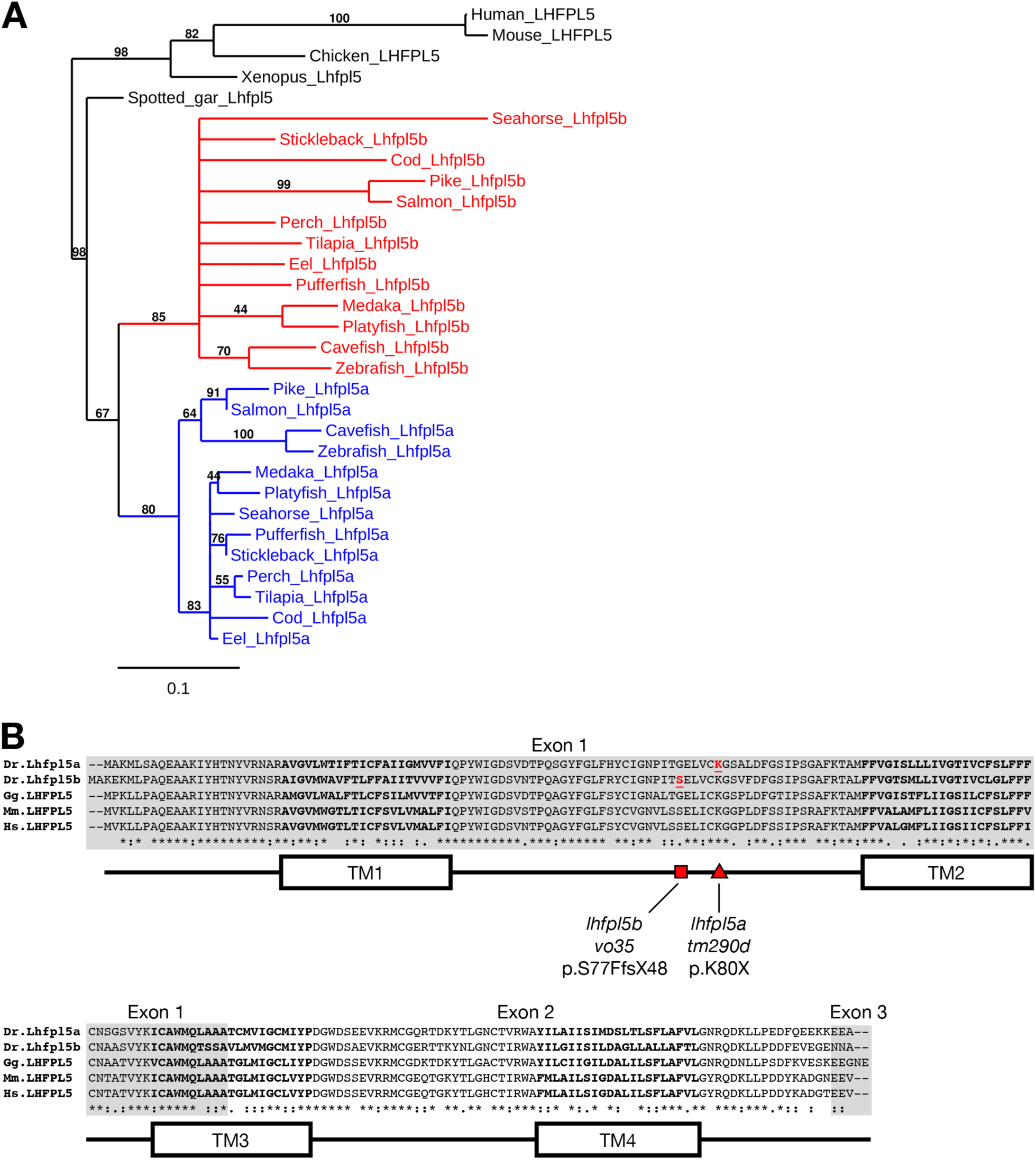
Duplicated *lhfpl5* genes in teleost fish. (**A**) Phylogenic tree of representative teleost and vertebrate Lhfpl5 protein sequences. See Supplemental Table 1 for species’ names and protein accession numbers. (**B**) Sequence alignment of Lhfpl5 proteins from zebrafish (*Danio rerio*, Dr), chicken (*Gallus gallus*, Gg), mouse (*Mus musculus*, Mm), and humans (*Homo sapiens*, Hs). Regions of the proteins coded for by exons 1 and 3 are shaded grey. Locations of the transmembrane (TM) helices 1-4 are shown in the linear protein structure diagram. The *lhfpl5a^tm290d^* and *lhfpl5b^vo35^* mutations are indicated by the red triangle and red square, respectively.

Alignment of the zebrafish Lhfpl5a and Lhfpl5b proteins with those from human, mouse and chicken reveals that both zebrafish ohnologs retain the same protein structure (Figure 1B). Zebrafish Lhfpl5a and Lhfpl5b are 76% identical and 86% similar to one another (Needleman-Wunsch alignment). Compared to human LHFPL5, Lhfpl5a and Lhfpl5b are 70 / 65 % identical and 86 / 81 % similar respectively. The ENU-generated mutation in *lhfpl5a* (*tm290d*) (Obholzer et al., 2012) is indicated by the red triangle (K80X). To investigate the function of *lhfpl5b*, we generated a Cas9-induced lesion in the *lhfpl5b* gene in a similar location as the *tm290d* mutation. We recovered a line with a 5 base pair deletion leading to a frameshift mutation (*lhfpl5b^vo35^*, red square; S77FfsX48).

### 4.2 *lhfpl5a* and *lhfpl5b* are expressed in distinct populations of sensory hair cells

To characterize the spatial and temporal patterns of *lhfpl5a/b* gene expression, we performed whole mount mRNA *in situ* hybridization on zebrafish larvae at 1, 2, and 5-days post-fertilization (dpf) (Figure 2). After one day of development, there are nascent hair cells of the presumptive anterior and posterior maculae in the developing ear, but no lateral line hair cells at this stage. We detect *lhfpl5a* expression in the presumptive anterior and posterior maculae at 1 dpf (Figure 2A). No signal for *lhfpl5b* was observed at this time point (Figure 2B). At 2 dpf, we observe a clear distinction in the expression patterns of the *lhfpl5a* and *lhfpl5b* genes (Figure 2C, D). *lhfpl5a* continues to be expressed in the ear, but expression is not observed in the newly deposited neuromasts. Conversely, we detect *lhfpl5b* expression exclusively in neuromasts at this stage, both on the head (Figure 2D) and trunk (data not shown). This divergence in *lhfpl5* ohnolog expression continues at 5 dpf, with *lhfpl5a* found exclusively in the sensory patches of the ear and *lhfpl5b* restricted to lateral line hair cells (Figure 2E-J). Taken together, our results suggest that both *lhfpl5a* and *lhfpl5b* have been retained since the teleost whole genome duplication because of their non-overlapping mRNA expression patterns.

**Figure 2.**
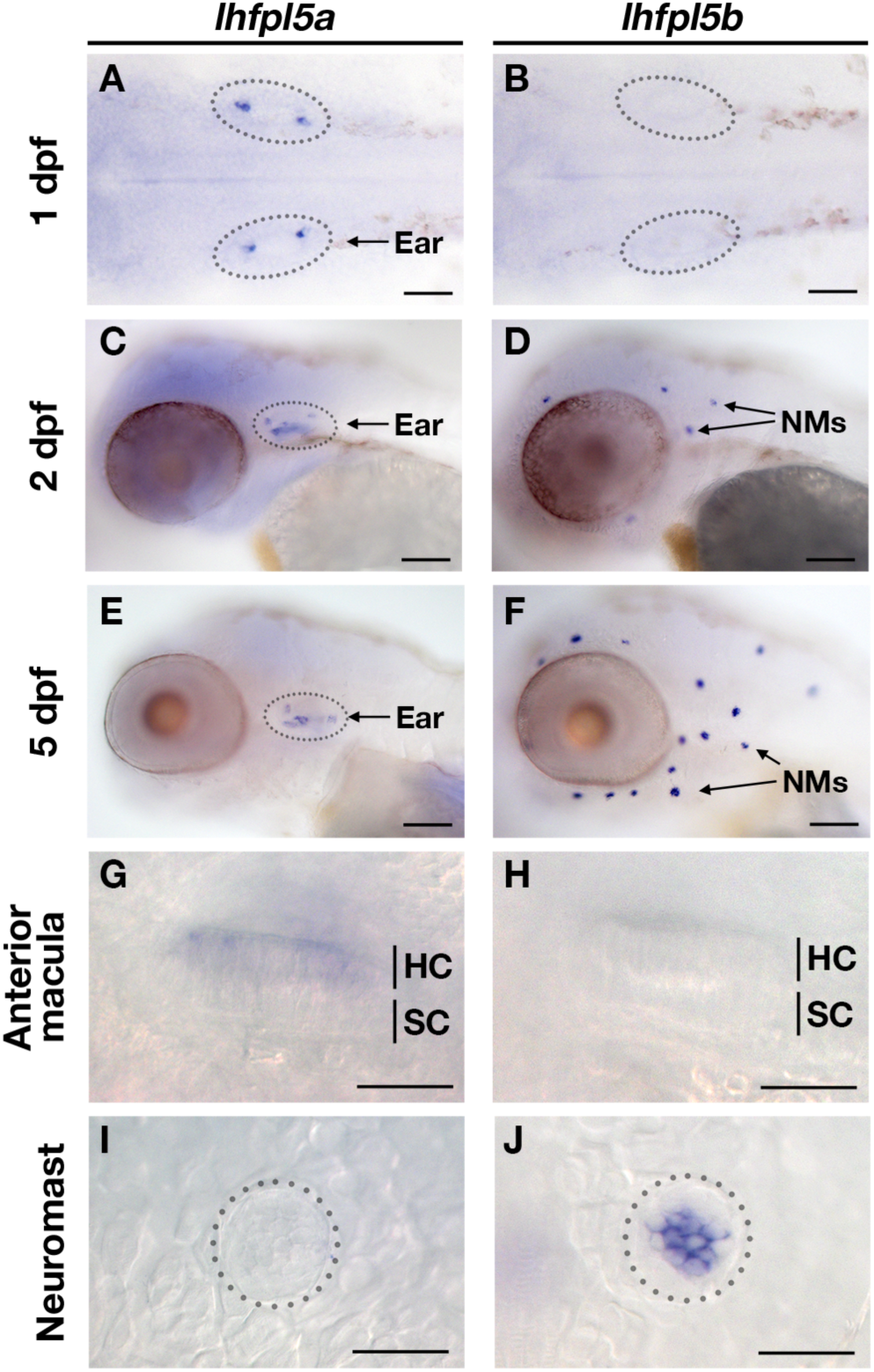
Zebrafish *lhfpl5a* and *lhfpl5b* genes are expressed in distinct populations of sensory hair cells. Whole mount mRNA *in situ* hybridization for *lhfpl5a* (**A, C, E, G, I**) and *lhfpl5b* (**B, D, F, H, J**) at 1, 2, and 5 days post-fertilization (dpf). Panels G - J are details from 5 dpf larvae. Abbreviations: NMs = neuromasts, HC = hair cell, SC = support cell. Scale bars: A, B = 50 µm; C - F = 100 µm; G - J = 25 µm.

### 4.3 *lhfpl5a* mediates mechanosensitivity of otic hair cells

*lhfpl5a^tm290d^* (*astronaut / asn*) mutants were initially characterized by balance defects, an absence of the acoustic startle reflex, and a lack of brainstem Ca^2+^ signals after acoustic startle. However, neuromast microphonic potentials were normal (Nicolson et al., 1998). In light of our *in situ* hybridization data (Figure 2), these results suggest that *lhfpl5a* is required for auditory and vestibular hair cell function only. To test this possibility, we performed several functional assays measuring the activity of hair cells in the otic capsule. First, we tested macular hair cell activity (Lu and DeSmidt, 2013; Yao et al., 2016) by recording extracellular microphonic potentials from the inner ears of *lhfpl5a^tm290d^* mutants, along with wild type and *lhfpl5b^vo35^* larvae at 3 dpf. We then measured baseline-to-peak amplitude of the first peak to quantify activity (Supplemental Figure 1A-D). Consistent with the initial characterization of the behavioral defects in *lhfpl5a^tm290d^* mutants, we did not detect robust microphonic potentials from the inner ear of *lhfpl5a^tm290d^* mutants (Figure 3A). We measured an average first peak microphonic of 53.1 µV from *lhfpl5a^tm290d^* mutants (n = 7), which was significantly less than the 296.2 µV average value of their WT siblings (n = 5; p = 0.01, Welch’s t-test). We believe that at least some of the signal detected in *lhfpl5a^tm290d^* mutants may be a stimulus artefact due the unusual rise time and similar observations we have made in other known transduction-null mutants.

**Figure 3.**
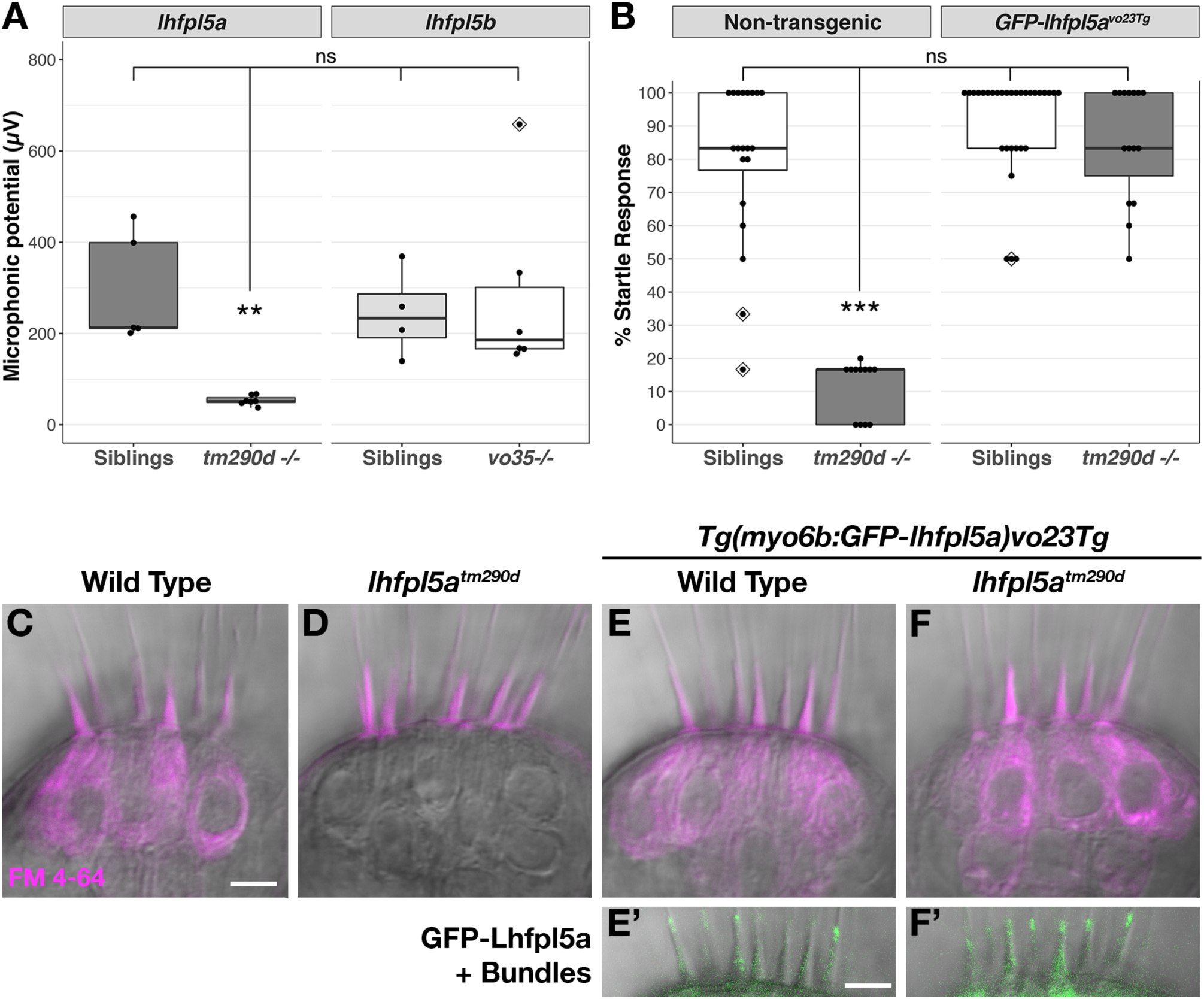
*lhfpl5a* is required for auditory and vestibular hair cell function. (**A**) Boxplot showing the first peak amplitude (µV) of microphonic recordings from the inner ear of 3 dpf wild type, *lhfpl5a^tm290d^*, and *lhfpl5b^vo35^* larvae. The boxes cover the inter-quartile range (IQR), and the whiskers represent the minimum and maximum datapoints within 1.5 times the IQR. Values from individual larvae are indicated by the black dots and outliers indicated by a diamond. Asterisks indicate p < 0.01 (**) by Welch’s t-test. ns = not significant. (**B**) Boxplot of the acoustic startle response in 6 dpf wild type and *lhfpl5a^tm290d^* mutants, with or without the *GFP-lhfpl5a vo23Tg* transgene. Values from individual larvae are indicated by the black dots and outliers indicated by a diamond. Asterisks indicate p < 0.001 (***) by Welch’s t-test. ns = not significant. (**C-F’**) FM 4-64 dye labeling assay for MET channel activity of inner ear hair cells from 6 dpf larvae (C - Wild type (*+/+ or +/-*), D - *lhfpl5a^tm290d^* mutants, E - Wild type *GFP-lhfpl5a vo23Tg*, F - GFP*-lhfpl5a vo23Tg; lhfpl5a^tm290^*). E’ and F’ are images of GFP-Lhfpl5a protein in the bundle of the hair cells shown above in panels E and F. n = 2, 3, 12, and 4 for the genotypes in panels C-F, respectively. Scale bars = 5 µm and apply to all images.

For the *lhfpl5b^vo35^* mutants (n = 6), we measured an average first peak microphonic potential of 280.9 µV. These values were not significantly different from their WT siblings (n = 4, average 243.8 µV; p = 0.7) but were significantly greater than the *lhfpl5a^tm290d^* mutants (p = 0.036). Thus, *lhfpl5a^tm290d^* mutants exhibit defects in inner ear function, while the *lhfpl5b^vo35^* mutation has no effect on these hair cells. To further confirm the lack of inner ear function in *lhfpl5a^tm290d^* mutants, we performed an acoustic startle test on larvae at 6 dpf (Figure 3B). Results confirm that *lhfpl5a^tm290d^* mutants are profoundly deaf, displaying little to no response to acoustic stimuli (p < 0.001 compared to all other genotypes). Finally, we examined the basal MET channel activity of hair cells of the inner ear by injecting FM 4-64 into the otic capsule and imaging the lateral cristae sensory patch. WT hair cells readily labeled with FM 4-64 whereas *lhfpl5a^tm290d^* mutant cells showed no sign of dye internalization (Figure 3C-D). These three tests confirmed that all sensory patches in the otic capsule are inactive in *lhfpl5a^tm290d^* mutants.

A *GFP-lhfpl5a* transgenic line of zebrafish - *Tg(myo6b:eGFP-lhfpl5a)vo23Tg* - has been reported previously (Erickson et al., 2017). GFP-Lhfpl5a is present at the tips of stereocilia, similar to the localization observed for mouse LHFPL5 (Mahendrasingam et al., 2017; Xiong et al., 2012). To demonstrate that the GFP-Lhfpl5a protein is functional, we assayed for rescue of the acoustic startle reflex and MET channel activity in homozygous *lhfpl5a^tm290d^* mutants expressing the transgene. The startle reflex of *lhfpl5a^tm290d^* mutants expressing GFP-Lhfpl5a were statistically indistinguishable from non-transgenic and transgenic WT siblings (p = 0.48 and 0.26 respectively; Figure 3B). Likewise, FM 4-64 dye-labeling in the inner ear was restored to *lhfpl5a^tm290d^* mutants expressing GFP-Lhfpl5a (Figure 3E-F’). From these results we conclude that the hair bundle-localized GFP-Lhfpl5a protein is functional and can rescue the behavioral and MET channel defects in *lhfpl5a^tm290d^* mutants.

### 4.4 *lhfpl5b* mediates mechanosensitivity of lateral line hair cells

The divergent expression patterns of the *lhfpl5* ohnologs suggests that *lhfpl5b* alone may be mediating mechanosensitivity of lateral line hair cells. To test this, we compared basal MET channel activity in neuromast hair cells of WT, *lhfpl5a^tm290d^* and *lhfpl5b^vo35^* larvae using an FM 1-43 dye uptake assay (Figure 4A-C). Strikingly, lateral line hair cells in *lhfpl5b^vo35^* mutants do not label with FM 1-43 while there is no difference in the intensity of FM 1-43 labeling of hair cells between WT and *lhfpl5a^tm290d^* mutants (Figure 4A-C, Supplemental Figure 2A - C). We quantified this loss of lateral line function in *lhfpl5b* mutants by measuring the average FM 1-43 fluorescence intensity per hair cell in each imaged neuromast of *lhfpl5b^vo35^* mutants and wild type siblings at 2 dpf (Figure 4D, E, H) and 5 dpf (Figure 4F-H). *lhfpl5b^vo35^* mutants exhibit a statistically significant decrease in FM 1-43 uptake at both 2 dpf and 5 dpf (Welch’s *t*-test, p < 0.001 both time points). While there is negligible FM dye labeling in the vast majority of *lhfpl5b^vo35^* mutant neuromasts, we occasionally observe labeling in some hair cells, most reproducibly in the SO3 neuromast. Occasional labeling persists in *lhfpl5a* / *lhfpl5b* double mutants, indicating that *lhfpl5a* is not partially compensating for the loss of *lhfpl5b* (Supplemental Figure 3A-D). The loss of MET channel activity in *lhfpl5b^vo35^* neuromasts is rescued by expression of the *GFP-lhfpl5a vo23Tg* transgene (Figure 4I, J; n = 7/7 mutant individuals). This result indicates that Lhfpl5a and Lhfpl5b are functionally interchangeable in this context.

**Figure 4.**
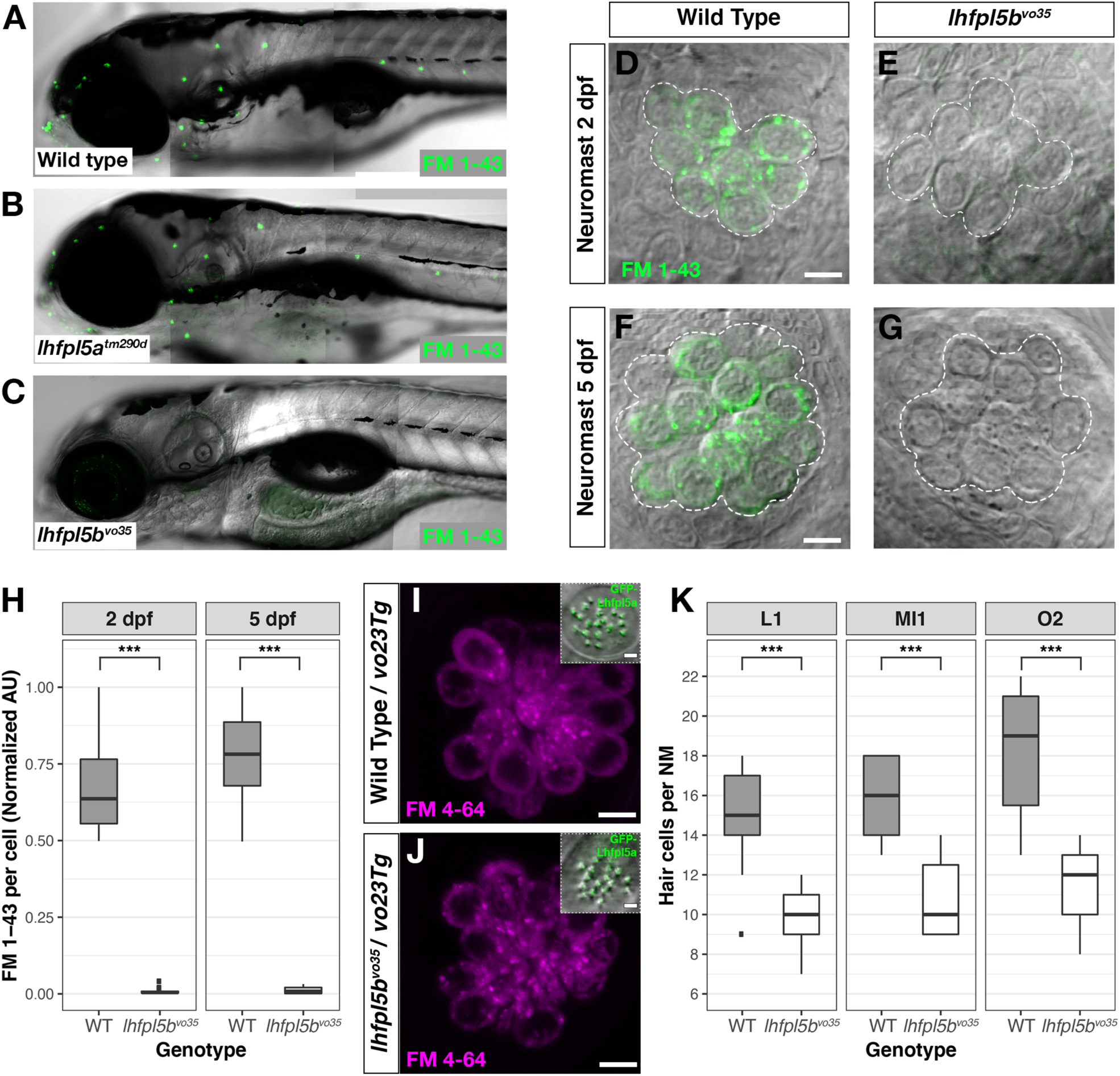
*lhfpl5b* is required for lateral line hair cell function. (**A-C**) Representative images of 5 dpf zebrafish larvae (A - Wild type; B - *lhfpl5a^tm290d^*; C - *lhfpl5b^vo35^*) labeled with the MET channel-permeant dye FM 1-43. (**D-G**) Representative images of individual neuromasts from 2 dpf and 5 dpf wild type and *lhfpl5b^vo35^* larvae labeled with FM 1-43. Dashed lines outline the cluster of hair cells in each neuromast. (**H**) Quantification of normalized FM 1-43 fluorescence intensity per hair cell of 2 dpf and 5 dpf neuromasts (n = 10 WT, 14 *lhfpl5b^vo35^* NMs at 2 dpf; n = 6 WT, 13 *lhfpl5b^vo35^* NMs each genotype at 5 dpf). The box plots cover the inter-quartile range (IQR), and the whiskers represent the minimum and maximum datapoints within 1.5 times the IQR. Asterisks indicate p < 0.001 (***) by Welch’s t-test. (**I, J**) Rescue of FM dye labeling in *lhfpl5b^vo35^* mutants (n = 7) by the *GFP-lhfpl5a* (*vo23Tg*) transgene. The GFP-Lhfpl5a bundle and FM 4–64 images are from the same NM for each genotype. (**K**) Quantification of hair cell number in L1, MI1, and O2 neuromasts from 5 dpf *lhfpl5b^vo35^* mutants (n = 11) and wild-type siblings (n = 11). The box plots are the same as in H. Asterisks indicate p < 0.001 (***) by Welch’s t-test. Scale bars = 5 µm, applies to panels D-G, I, J; 2 µm in I, J insets).

We have previously reported a reduced number of neuromast hair cells in other transduction mutants such as *myo7aa (mariner)*, *pcdh15a (orbiter)*, and *tomt (mercury)* (Erickson et al., 2017; Seiler et al., 2005). The ultimate cause for this reduction in hair cell number is not known. If *lhfpl5b^vo35^* mutants are defective in mechanotransduction, we would expect a similar decrease in the number of neuromast hair cells. To test this idea, we compared hair cell counts between wild type, *lhfpl5a^tm290d^*, and *lhfpl5b^vo35^* larvae at 5 dpf. As expected, wild type and *lhfpl5a^tm290d^* mutants have statistically equivalent numbers of hair cells in the neuromasts surveyed (Supplemental Figure 2D, p = 0.58). In *lhfpl5b^vo35^* mutant neuromasts, we observe a statistically significant decrease in the number of hair cells (Figure 4K, n = 11 individuals per genotype, 3 NM each, p < 0.001 each NM type by Welch’s *t*-test; representative neuromasts are shown in Supplemental Figure 4). This decrease in the number of neuromast hair cells is consistent with *lhfpl5b* mutants being deficient in mechanotransduction in this cell type.

### 4.5 *lhfpl5* is not required for Tmc localization to the hair bundle in zebrafish hair cells

Previous studies have demonstrated that loss of LHFPL5 leads to a ∼90% reduction in the peak amplitude of the transduction current in mouse cochlear outer hair cells (Xiong et al., 2012). Single channel recordings from *Lhfpl5* mutants revealed that the remaining transduction current in *Lhfpl5* mutants was mediated by TMC2 (Beurg et al., 2015). Using antibodies to detect endogenous protein, TMC1 was found to be absent from the bundle of *Lhfpl5* mouse mutants. However, injectoporated Myc-TMC2 was still able to localize to the hair bundle, supporting the idea that LHFPL5 is required for the targeting of TMC1, but not TMC2. The localization of exogenously-expressed TMC1 in *Lhfpl5* mutants was not reported.

To determine if the Tmcs require Lhfpl5a for localization to the hair bundle of zebrafish hair cells, we bred stable transgenic lines expressing GFP-tagged versions of Tmc1 (*vo27Tg*) and Tmc2b (*vo28Tg*) (Erickson et al., 2017) into the *lhfpl5a^tm290d^* mutant background and imaged Tmc localization in the lateral cristae of the ear. In contrast to what was observed in mouse cochlear hair cells, both Tmc1-GFP and Tmc2b-GFP are still targeted to the hair bundle in *lhfpl5a^tm290d^* mutants (n = 11/11 and 9/9 individuals respectively; Figure 5A-D’). We also observe Tmc2b-GFP localization in the neuromast hair bundles of *lhfpl5b^vo35^* mutants (n = 7/7 individuals; Figure 5E-F’). Additionally, we do not observe any rescue of hair cell function in *lhfpl5b* mutants expressing the Tmc2b-GFP protein (Supplemental Figure 5A-C). As such, these results suggest that GFP-tagged Tmc1 and Tmc2b do not require Lhfpl5 for hair bundle localization in zebrafish hair cells.

**Figure 5.**
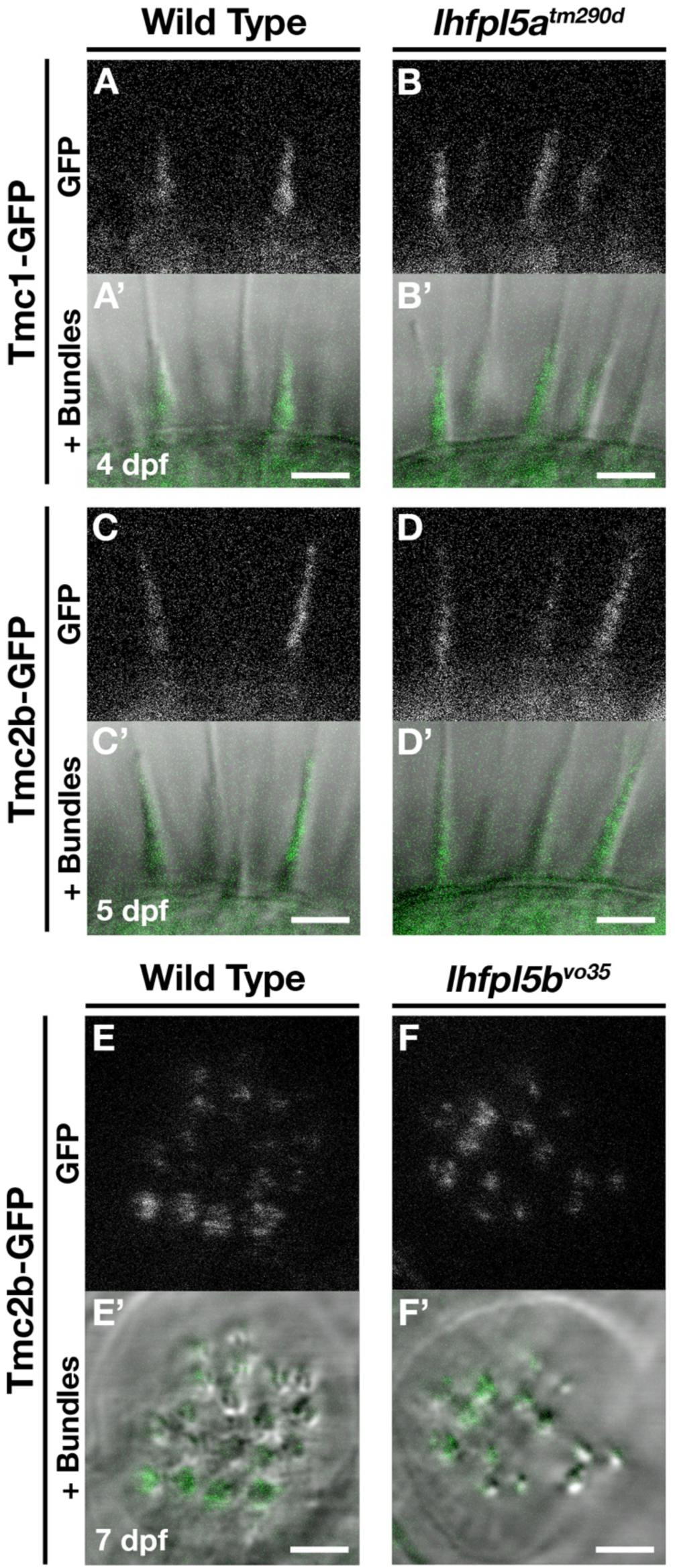
Tmc proteins do not require Lhfpl5a or Lhfpl5b for localization to the stereocilia of zebrafish hair cells. (**A – D’**) Representative images of Tmc1-GFP (*vo27Tg*) or Tmc2b-GFP (*vo28Tg*) in the lateral cristae of wild type (**A, A’, C, C’**) and *lhfpl5a^tm290d^* (**B, B’, D, D’**) larvae. The GFP-only channel is shown in panels A - D and overlaid with a light image of the bundles in A’ - D’. (**E – F’**) Representative images of Tmc2b-GFP (*vo28Tg*) in the neuromasts of wild type (**E, E’**) and *lhfpl5b^vo35^* (**F, F’**) larvae. The GFP-only channel is shown in panels E and F and overlaid with a light image of the bundles in E’ and F’. Scale bars = 3 µm in all panels.

### 4.6 GFP-Lhfpl5a localization in stereocilia requires Pcdh15a, Cdh23, and Myo7aa

Previous work has shown that PCDH15 and LHFPL5 are mutually dependent on one another to correctly localize to the site of mechanotransduction in mouse cochlear hair cells (Mahendrasingam et al., 2017; Xiong et al., 2012). Likewise in zebrafish, Lhfpl5a is required for proper Pcdh15a localization in the hair bundle, as determined Pcdh15a immunostaining and the expression of a Pcdh15a-GFP transgene in *lhfpl5a^tm290d^* mutants (Maeda et al., 2017). Using the previously characterized antibody against Pcdh15a, we observe the highest level of Pcdh15a staining at the apical part of the hair bundle, with less intense punctate staining throughout the stereocilia in 3 dpf wild type larvae (Figure 6A). *lhfpl5a^tm290d^* mutants exhibit splayed hair bundles, with low levels of Pcdh15a staining restricted to the tip of each stereocilium (n = 5/5 individuals; Figure 6B), confirming our previous report.

**Figure 6.**
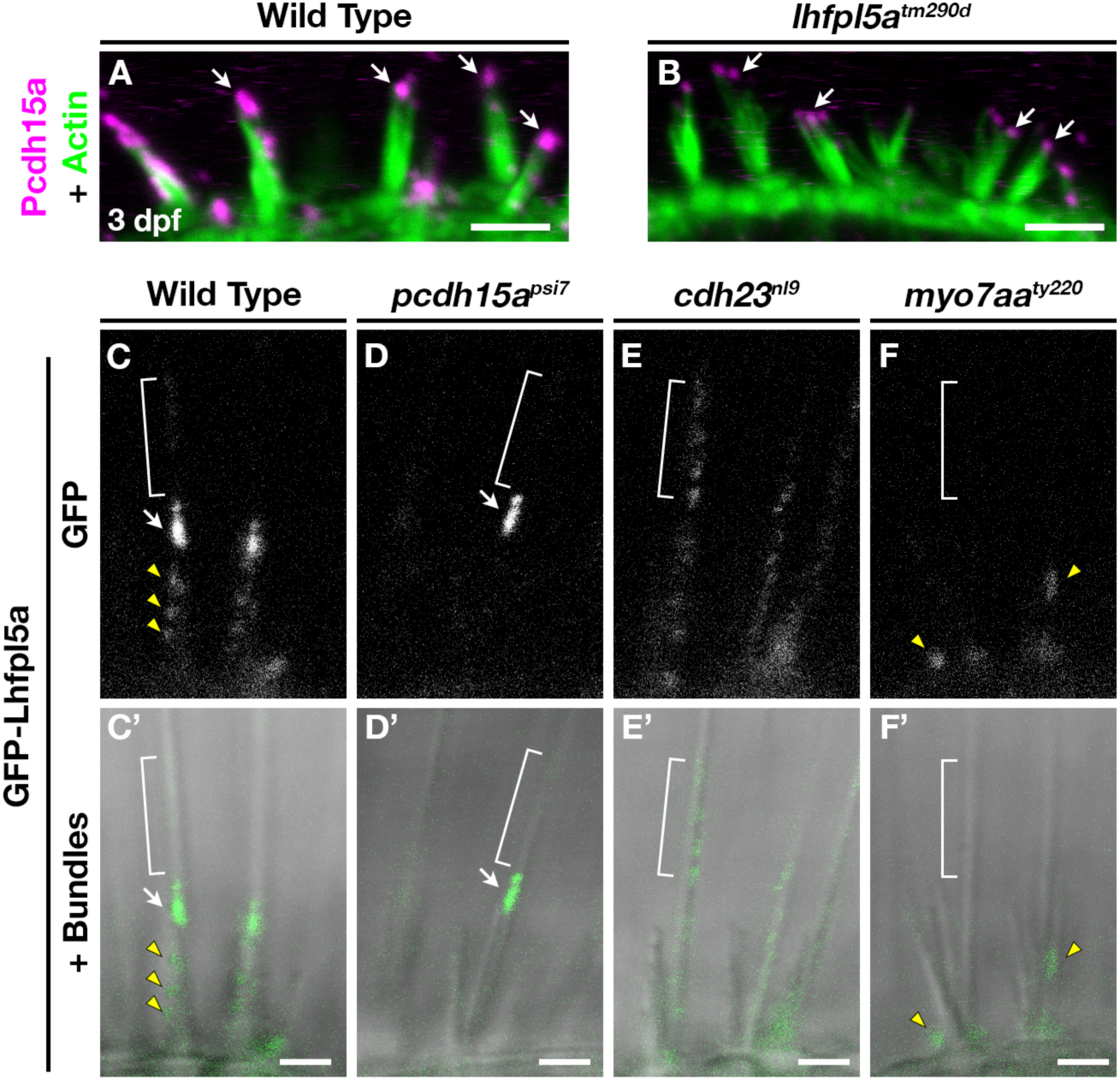
Lhfpl5a requires MET complex proteins Pcdh15a, Cdh23, and Myo7a for normal localization in the stereocilia of zebrafish hair cells. (**A, B**) Immunostain of Pcdh15a (magenta) in the lateral cristae of wild type and *lhfpl5a^tm290d^* mutants at 3 dpf. Phalloidin-stained actin of the hair bundle is shown in green. Arrows indicate areas of Pcdh15a accumulation. (**C-F’**) Representative images of GFP-Lhfpl5a (*vo23Tg*) in the lateral cristae hair bundles of wild type (**C, C’**) and *pcdh15a^psi7^* (**D, D’**), *cdh23^nl9^* (**E, E’**), and *myo7aa^ty220^* (**F, F’**) mutants. White arrows indicate GFP signal in the presumptive kinocilial linkages, yellow arrow heads indicate GFP signal in the stereocilia or the base of the hair bundle, and brackets indicate GFP signal in the kinocilium. Scale bars = 3 µm in A, B; 2 µm in C-F’.

To determine if Lhfpl5a also depends on Pcdh15a for its localization, we imaged GFP-Lhfpl5 in the lateral cristae of *pcdh15a^psi7^* homozygous mutants. In wild type larvae, GFP-Lhfpl5a is distributed throughout the hair bundle at the tips of stereocilia (yellow arrow heads), with a higher intensity of GFP evident in the tallest rows, possibly at the sites of kinociliary links (Figure 6C, C’, arrow). In *pcdh15a^psi7^* mutants, GFP-Lhfpl5a is absent from the hair bundle except for the tallest rows of stereocilia adjacent to the kinocilium (n = 4 individuals; Figure 6D, D’, arrow). We observed similar results using the *pcdh15a^th263b^* allele (n = 6/6 individuals; Supplemental Figure 6A, B). Consistent with our earlier studies (Maeda et al., 2017; Seiler et al., 2005), we observe splayed hair bundles in both *pcdh15a* alleles suggesting a loss of connections and tip links between the stereocilia. These results indicate that GFP-Lhfpl5a requires Pcdh15a for stable targeting to the shorter stereocilia, but Pcdh15a is not required for retaining GFP-Lhfpl5a in the tallest rows of stereocilia adjacent to the kinocilium.

The observation that some GFP-Lhfpl5a signal remains in the tallest stereocilia of *pcdh15a* mutants suggests that additional proteins are involved in targeting Lhfpl5a to these sites in zebrafish vestibular hair cells. Previous research has shown that mouse CDH23 still localizes to tips of stereocilia in PCDH15-deficient mice (Boëda et al., 2002; Senften et al., 2006) and that CDH23 is a component of kinociliary links (Goodyear et al., 2010; Siemens et al., 2004). Based on these reports, we tested whether Cdh23 was required for GFP-Lhfpl5a localization. In *cdh23^nl9^* mutants, GFP-Lhfpl5 is sparsely distributed throughout the length of the kinocilium (n = 4/4 individuals, Figure 6E, E’). We also observe a redistribution of GFP-Lhfpl5a into the kinocilium using the *cdh23^tj264^* allele (n = 4/4 individuals; Supplemental Figure 6C). In both *cdh23* alleles, we see occasional GFP signal in the stereocilia as well, though this localization is not as robust. Interestingly, these results contrast with those from mouse cochlear hair cells which showed that Lhfpl5 does not require Cdh23 for bundle localization (Zhao et al., 2014). The same study also could not detect any biochemical interaction between LHFPL5 and proteins of the upper tip link density including CDH23, USH1C / Harmonin, or USH1G / Sans. Together, our data indicate an unexpected role for Cdh23 in the localization of GFP-Lhfpl5a to the region of kinocilial links in zebrafish vestibular hair cells.

Myosin 7A (MYO7A) is an actin-based motor protein required for the localization of many proteins of the hair bundle, including PCDH15 and USH1C / Harmonin, along with several proteins of the Usher type 2 complex (Boëda et al., 2002; Lefevre et al., 2008; Maeda et al., 2017; Morgan et al., 2016; Senften et al., 2006; Zou et al., 2017). Results in both mice and zebrafish suggest that MYO7A is not required for CDH23 bundle localization (Blanco-Sanchez et al., 2014; Senften et al., 2006). However, its role in Lhfpl5 localization has not been examined. In *myo7aa^ty220^* (*mariner*) mutant hair cells, GFP-Lhfpl5a localization in hair bundle is severely disrupted (n = 5/5 individuals; Figure 6 F, F’). We observe GFP signal near the base of hair bundles and occasionally in stereocilia (yellow arrowheads), but do not see robust Lhfpl5a localization in the presumptive kinocilial links nor to the kinocilium itself. Taken together, our results using the GFP-Lhfpl5a transgene suggest that Pcdh15a, Cdh23, and Myo7aa all play distinct roles in Lhfpl5 localization in the bundle of zebrafish vestibular hair cells.

## 5 Discussion

In this study, we provide support for the following points: 1) The ancestral *lhfpl5* gene was likely duplicated as a result of the teleost whole genome duplication, leading to the *lhfpl5a* and *lhfpl5b* ohnologs described in this paper. 2) In zebrafish, there has been subfunctionalization of these genes as a result of their divergent expression patterns. As determined by mRNA *in situ* hybridization, *lhfpl5a* is expressed only in auditory and vestibular hair cells of the ear while *lhfpl5b* is expressed solely in hair cells of the lateral line organ. Consistent with their expression patterns, we show that each ohnolog mediates mechanotransduction in the corresponding populations of sensory hair cells in zebrafish. 3) Targeting of GFP-tagged Tmcs to the hair bundle is independent of Lhfpl5a function. 4) Proper targeting of GFP-tagged Lhfpl5a to the hair bundle requires the tip link protein Pcdh15a, but as in mice, Lhfpl5a can localize to regions of the hair bundle independently of Pcdh15 function. Additionally, we demonstrate novel requirements for Cdh23 and the Myo7aa motor protein in Lhfpl5 localization.

### 5.1 Duplicated zebrafish genes in hair cell function

With regards to genes involved in hair cell function, the zebrafish duplicates of *calcium channel, voltage-dependent, L type, alpha 1D* (*cacna1d* / *cav1.3*), *C-terminal binding protein 2* (*ctbp2* / *ribeye*)*, myosin 6* (*myo6*)*, otoferlin* (*otof*), *pcdh15*, and *tmc2* have been analyzed genetically. In the cases of *cacna1db*, *pcdh15b* and *myo6a*, these ohnologs are no longer functional in hair cells (Seiler et al., 2004, 2005; Sidi et al., 2004). For the *ctbp2*, *otoferlin*, and *tmc2* duplicates, both genes are required in at least partially overlapping populations of hair cells (Chou et al., 2017; Lv et al., 2016; Maeda et al., 2014; Sheets et al., 2011). In contrast, our results for the *lhfpl5* ohnologs suggest that each gene is expressed and functionally required in distinct, non-overlapping hair cell lineages. To our knowledge, this is the first description of duplicated genes whose expression patterns have cleanly partitioned between inner ear and lateral line hair cells.

### 5.2 Subfunctionalization of zebrafish *lhfpl5* ohnologs

We constructed a phylogenetic tree using publicly available Lhfpl5 protein sequence data (Figure 1) Our results suggest that the ancestral *lhfpl5* gene was duplicated during the teleost WGD event and that both genes have been retained throughout the teleost lineage (Figure 1). Our conclusion that *lhfpl5a* and *lhfpl5b* are ohnologs is substantiated by a recent study of ohnologs in teleosts (Singh and Isambert, 2019). In all four teleost species surveyed, Singh and Isambert show that *lhfpl5a* and *lhfpl5b* are true ohnologs that arose from the teleost-specific WGD under the strictest criteria used in their study.

Some kind of selection pressure is required for the retention of both gene ohnologs (Glasauer and Neuhauss, 2014). Our in situ hybridization results show that *lhfpl5a* and *lhfpl5b* are expressed in distinct populations of hair cells and support the idea that the zebrafish *lhfpl5* ohnologs were retained because of their divergent expression patterns (Figure 2). Analysis of hair cell function in the different *lhfpl5* mutants agrees with the gene expression results, showing that *lhfpl5a* is required for transduction in the ear while *lhfpl5b* plays the same role in the lateral line organs. However, rescue of the MET channel defects in *lhfpl5b^vo35^* mutants by GFP-tagged Lhfpl5a suggests that the functions of these ohnologs are at least partially interchangeable. This result is not surprising given the high degree of similarity between the Lhfpl5a and 5b proteins. As such, the subfunctionalization of the *lhfpl5* ohnologs appears to be caused by the divergence in their expression patterns rather than functional differences in their protein products. It is possible that a similar mechanism is responsible for the retention of both genes in other teleost species as well, though the expression patterns of the *lhfpl5* ohnologs have not been examined in other fish.

The non-overlapping expression of the *lhfpl5* ohnologs provides us with a unique genetic tool to study lateral line function. Most mechanotransduction mutants in zebrafish disrupt both inner ear and lateral line hair cell function (Nicolson, 2017). As such, we are unable to assess the role of the lateral line independently of auditory and vestibular defects, which are lethal for larval fish. The *lhfpl5b* mutants are adult viable (data not shown) and represent a possible genetic model for understanding the lateral line in both larval and adult fish. Future studies on adults will determine whether the lateral line remains non-functional and whether *lhfpl5b* contributes to inner ear function in the mature auditory and vestibular systems.

### 5.3 The molecular requirements for bundle localization of MET complex proteins differs between mouse and zebrafish hair cells

Our understanding of mechanotransduction at the molecular level is heavily informed by work done using mouse cochlear hair cells. However, analysis of zebrafish inner ear and lateral line hair cells can lead to a more complete picture of how vertebrate sensory hair cells form the MET complex. At their core, the genetic and molecular bases for mechanotransduction are well conserved between mouse and zebrafish hair cells. For example, the tip link proteins Cdh23 and Pcdh15a, MET channel subunits Tmc1/2, Tmie, and Lhfpl5, along with additional factors such as Tomt, Cib2, and Myo7aa are all necessary for MET channel function in both mice and fish (Cunningham et al., 2017; Erickson et al., 2017; Giese et al., 2017; Gleason et al., 2009; Kawashima et al., 2011; Nicolson et al., 1998; Pacentine and Nicolson, 2019; Senften et al., 2006; Siemens et al., 2004; Söllner et al., 2004; Xiong et al., 2012; Zhao et al., 2014). However, while each of these factors are required for mechanotransduction in vertebrate hair cells, the details of how they contribute to MET channel function may differ depending on the particular type of hair cell or the vertebrate species. For example, Tmie is required for Tmc localization to the hair bundle in zebrafish (Pacentine and Nicolson, 2019), but this does not appear to be the case in mouse cochlear hair cells (Zhao et al., 2014). The evolutionary pressures that lead to these differences between mouse and zebrafish hair cells are not understood.

Based on multiple lines of evidence gathered from mice, TMC1 and TMC2 have different requirements for LHFPL5 regarding localization to the hair bundle (Beurg et al., 2015). Endogenous TMC1 is absent from the bundle in LHFPL5-deficient cochlear hair cells with the remaining MET current mediated through TMC2. Exogenously-expressed TMC2 is still targeted correctly in *Lhfpl5* mutants, but it was not reported whether the same is true for exogenous TMC1, nor whether similar requirements hold for vestibular hair cells. As such, the role of LHFPL5 in mechanotransduction is not clear: is LHFPL5 required primarily for targeting TMC1 to the hair bundle, or does LHFPL5 mediate MET channel function in other ways?

For our study, we used stable transgenic lines expressing GFP-tagged Tmc1 and Tmc2b to show that the Tmcs do not require Lhfpl5a for targeting to the stereocilia of zebrafish vestibular hair cells. Nor is Lhfpl5b required for Tmc2b-GFP localization in the hair bundle of neuromast hair cells. The occasional FM labeled-hair cell in *lhfpl5b* and *lhfpl5a/5b* mutant neuromasts is consistent with the continued ability of the Tmcs to localize to the hair bundle. This observation is reminiscent of the sporadic, low level of FM dye labeling in the cochlear hair cells of *Lhfpl5* mutant mice (György et al., 2017). Partial compensation by yet another member of the *lhfpl* family remains as a possible explanation for the remaining basal channel function. Lastly, we find that exogenous Tmc expression does not rescue MET channel activity in either *lhfpl5* mutant zebrafish. Taken together, our results support a Tmc-independent role for Lhfpl5 proteins in mechanotransduction.

What then is the role of LHFPL5 in vertebrate mechanotransduction? LHFPL5 and PCDH15 are known to form a protein complex and regulate each other’s localization to the site of mechanotransduction in mouse cochlear hair cells (Ge et al., 2018; Mahendrasingam et al., 2017; Xiong et al., 2012). However, their localization to the hair bundle is not completely dependent on one another. For example, tips links are not completely lost in *Lhfpl5^-/-^* cochlear hair cells and exogenous overexpression of PCDH15 partially rescues the transduction defects in *Lhfpl5* mutant mice (Xiong et al., 2012). Similarly, a detailed immunogold study in wild type and *Pcdh15^-/-^* cochlear hair cells showed that there are PCDH15-dependent and PCDH15-independent sites of LHFPL5 localization (Mahendrasingam et al., 2017). PCDH15 is required for stable LHFPL5 localization at the tips of ranked stereocilia in association with the MET channel complex. However, LHFPL5 is still targeted to shaft and ankle links, unranked stereocilia, and the kinocilium in P0-P3 inner and outer hair cells from both wild type and *Pcdh15^-/-^* mutant mice (Mahendrasingam et al., 2017). Whether the kinocilial localization would remain in mature cochlear hair cells is not known because the kinocilia degenerate by P8 in mice and detailed LHFPL5 localization in vestibular hair cells (which retain their kinocilium) has not been reported.

Zebrafish hair cells retain a kinocilium throughout their life, thus providing a different context in which to examine Lhfpl5 localization. In wild type hair cells, GFP-Lhfpl5a is present at the tips of the shorter ranked stereocilia, but the most robust signal is detected in the tallest stereocilia adjacent to the kinocilium. In *pcdh15a* mutants, only this “kinocilial link”-like signal remains in the hair bundle. This result is similar to the LHFPL5 localization reported at the tips of the tallest stereocilia of P3 cochlear hair cells from wild type mice, a region where LHFPL5 would presumably not be associated with either PCDH15 or the TMCs (Mahendrasingam et al., 2017). Since the kinocilium does not degenerate in zebrafish hair cells, our results suggest that Lhfpl5a is normally targeted to these kinocilial links in wild type cells and that its retention at this site does not require Pcdh15a. Thus, there are PCDH15-dependent and PCDH15-independent mechanisms of LHFPL5 localization in both mouse and zebrafish hair cells. These results suggest that LHFPL5 performs as-of-yet uncharacterized PCDH15-independent functions as well.

Cdh23 is thought to form part of the linkages between the kinocilium and adjacent stereocilia, and therefore may play a role in retaining Lhfpl5a at these sites. Consistent with this hypothesis, *cdh23* mutants do not exhibit GFP-Lhfpl5a accumulation in stereocilia. Instead, GFP-Lhfpl5a is diffusely distributed throughout the kinocilium. These data suggest that normal Lhfpl5a localization requires both Pcdh15a and Cdh23 function, albeit in distinct ways. This result differs from previous reports which found that Lhfpl5 still localized to the tips of stereocilia in CDH23-deficient cochlear hair cells of mice. The same study also could not detect a biochemical interaction between Lhfpl5 and CDH23 via co-IP in heterologous cells (Xiong et al., 2012). Given these contrasting results for mouse and zebrafish hair cells, it seems that the requirement for Cdh23 in the localization of LHFPL5 is not a universal one.

Given the available biochemical evidence, it is possible that the association between Lhfpl5a and Cdh23 is indirect through an as-of-yet unidentified protein or protein complex. Based on the defects in GFP-Lhfpl5a localization in *myo7aa* mutants, Myo7aa is a candidate member of this uncharacterized protein complex. Taken together, these results highlight the fact that, although all sensory hair cells share many core genetic and biochemical features, there are important details that differ between the various types of hair cells and between vertebrate species. Analyzing these differences will allow for a more comprehensive understanding of vertebrate hair cell function and the underlying principles of mechanotransduction.

## 7 Author Contributions

TE and TN conceived and designed the study; TE, IVP, AV, and RC collected and analyzed the data, TE wrote the manuscript with Methods sections contributed by IVP and AV and editorial input from TN and IVP.

## 9 Acknowledgments

We thank Eliot Smith for assistance with CRISPR-Cas9 knockout of *lhfpl5b*. We also thank Leah Snyder, Matthew Esqueda, and members of the Erickson lab for their assistance with animal husbandry.

## 11 Figures

**Supplemental Table 1.**
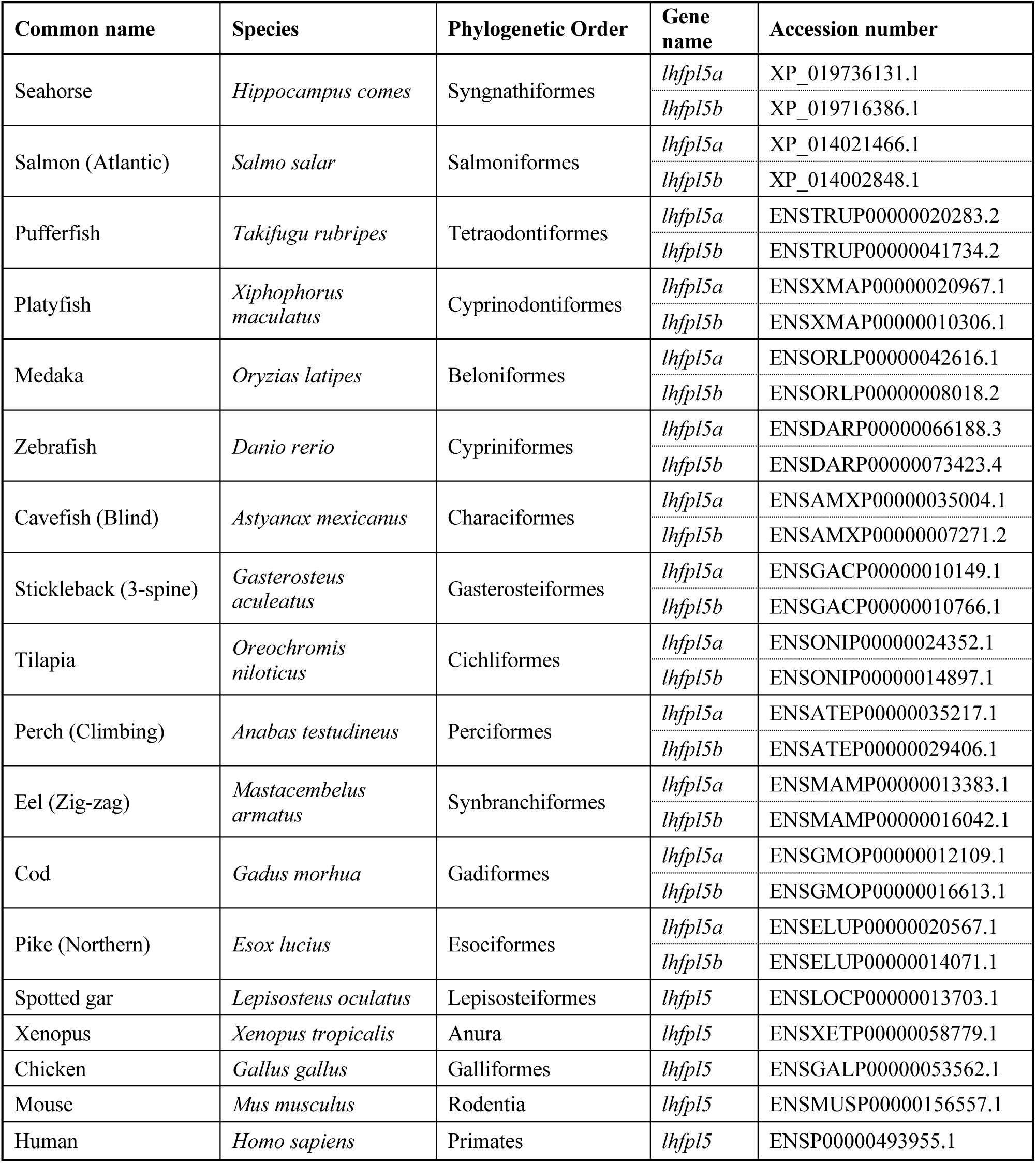
Lhfpl5 proteins used to construct the phylogenetic tree in Figure 1A.

**Supplemental Figure 1.**
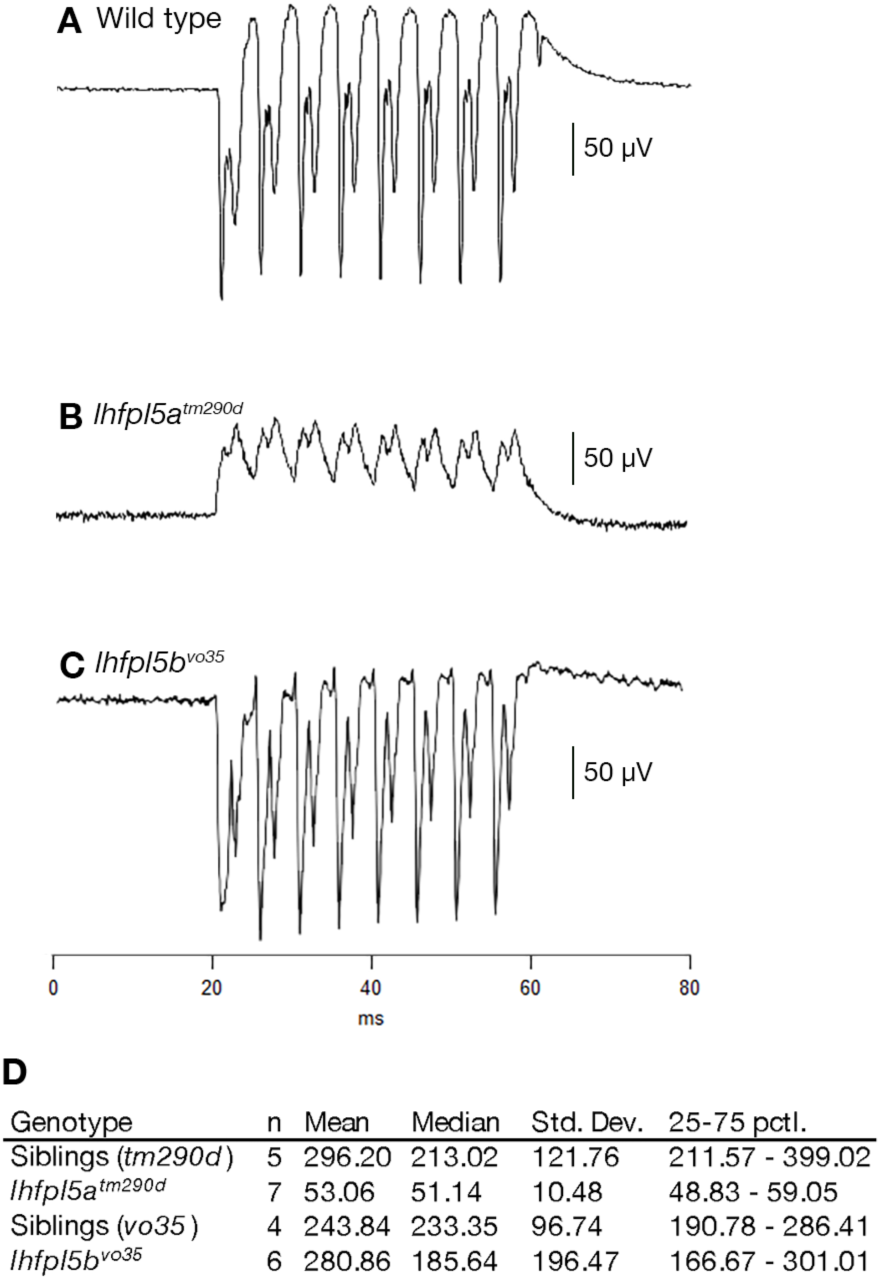
Representative microphonic traces from the inner ears of 3 day-old wild type (**A**), *lhfpl5a^tm290d^* (**B**), and *lhfpl5b^vo35^* (**C**) larvae. **D** – Table of the n-values, mean, median, standard deviation, and 25 - 75 percentile values for the first peak microphonic values graphed in Figure 3A.

**Supplemental Figure 2.**
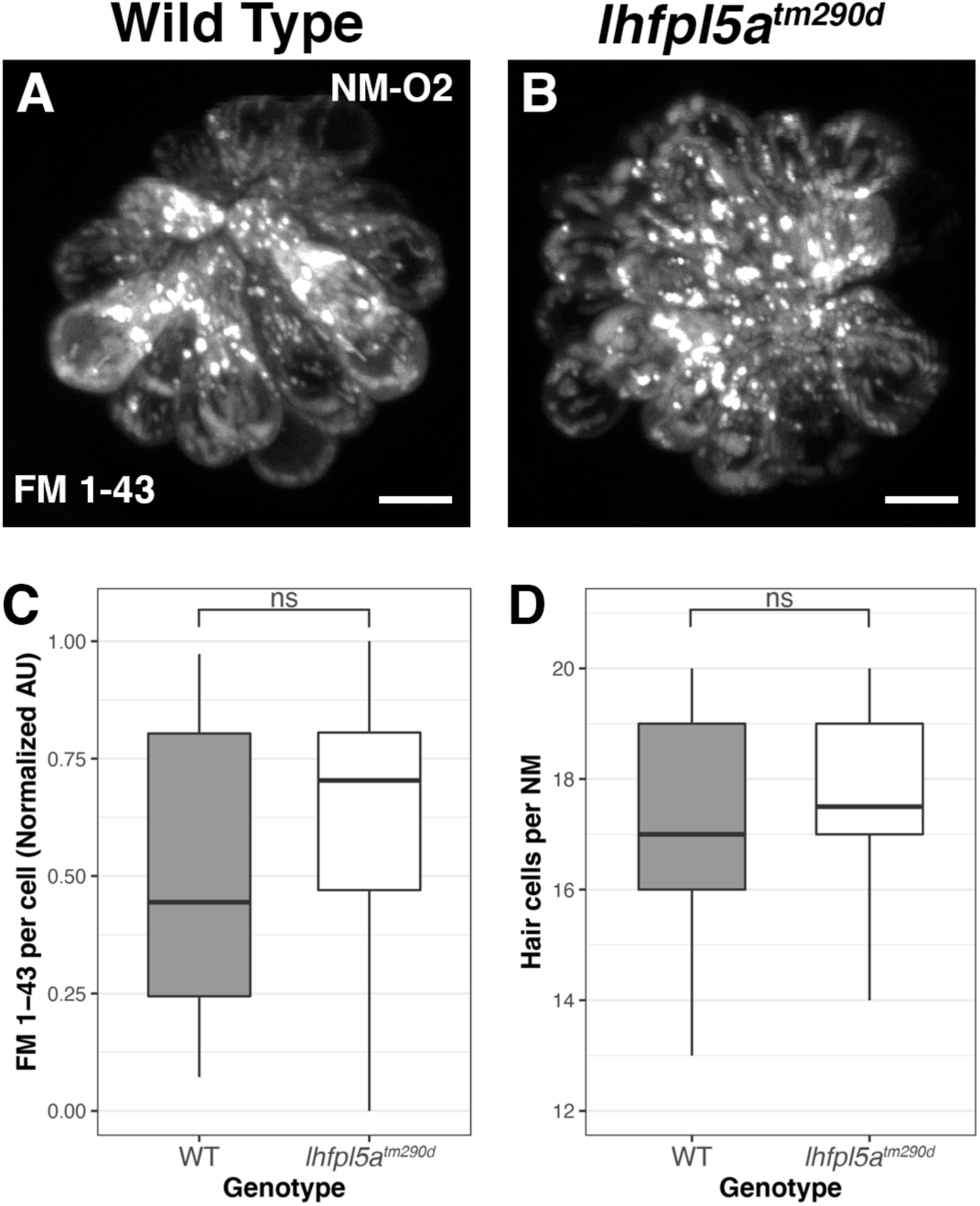
Comparison of basal MET channel activity in neuromasts from wild type and *lhfpl5a^tm290d^* mutant larvae. (**A, B**) Representative images of FM 1-43 fluorescence in lateral line neuromasts from wild type (A) and *lhfpl5a^tm290d^* (B) mutants at 5 dpf. (**C**) Quantification of normalized FM 1-43 fluorescence intensity per hair cell in 5 dpf neuromasts (n = 6 WT, 6 *lhfpl5a^tm290d^* larvae, 3 NMs per larvae). The box plots cover the inter-quartile range (IQR), and the whiskers represent the minimum and maximum datapoints within 1.5 times the IQR. p = 0.2229; ns = not significant. (**D**) Quantification of hair cell number in neuromasts from 5 dpf *lhfpl5a^tm290d^* mutants and wild-type siblings, as determined by counting FM-positive hair cells. The same larvae and neuromasts were used as in C. p = 0.5828; ns = not significant by Welch’s t-test.

**Supplemental Figure 3.**
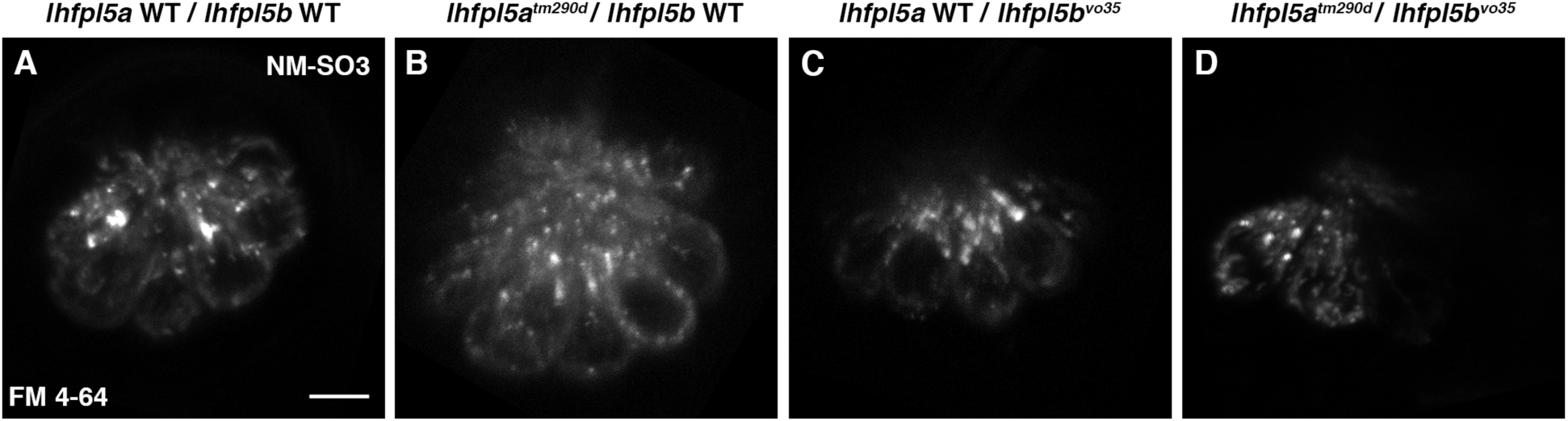
Examples of residual basal MET channel activity in the SO3 neuromast hair cells of *lhfpl5b^vo35^* mutants and *lhfpl5a^tm290d^*; *lhfpl5b^vo35^* double mutants. FM 4-64 labeled SO3 neuromasts from wild type (**A**), *lhfpl5a^tm290d^* (**B**), *lhfpl5b^vo35^* (**C**), and *lhfpl5a^tm290d^*; *lhfpl5b^vo35^* (**D**) larvae at 5 dpf. Scale bar = 5 µm, applies to all panels.

**Supplemental Figure 4.**
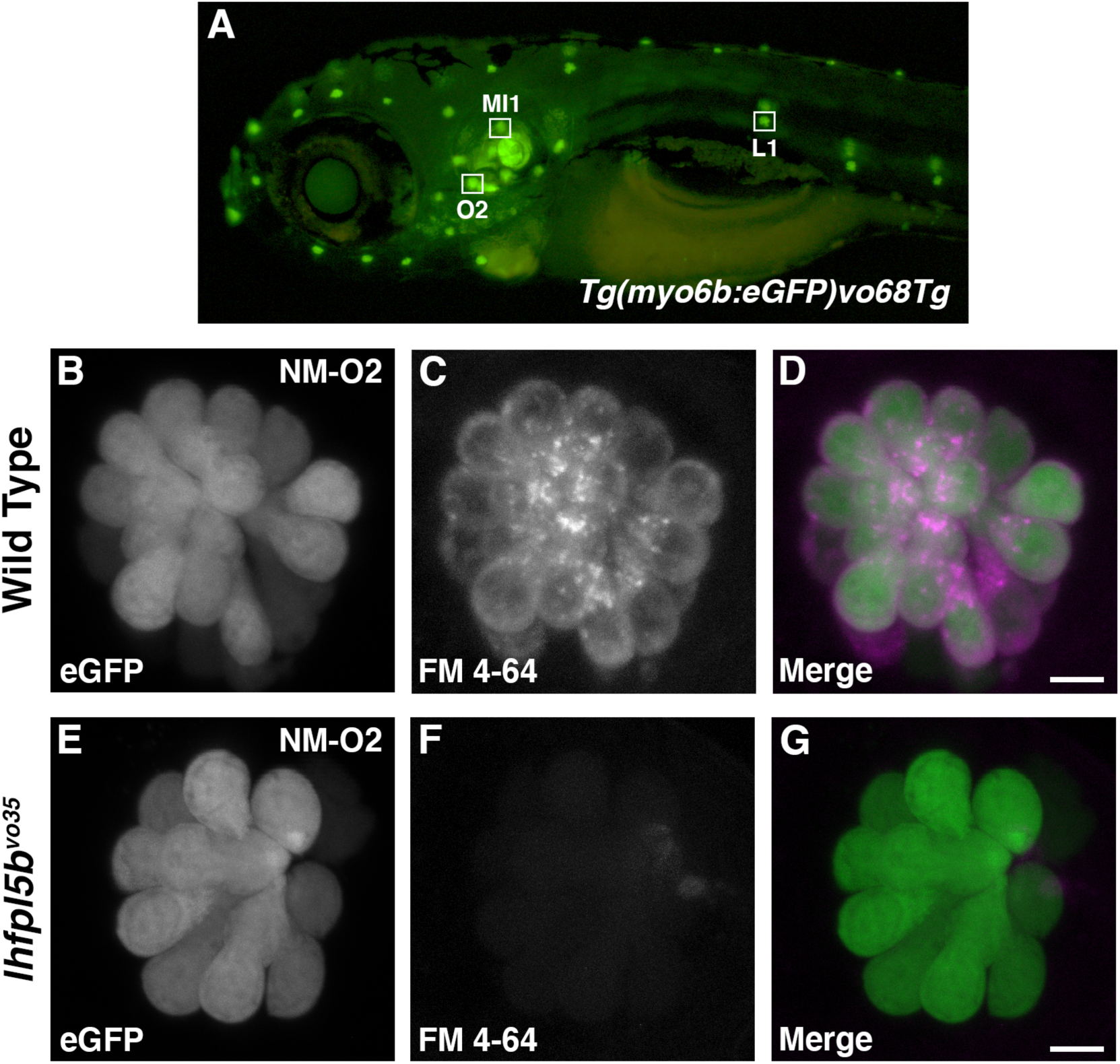
(**A**) *Tg(myo6b:GFP)vo68Tg* larvae at 5 dpf. The neuromasts used for hair cell counting are labeled. (**B – G**) Representative images of GFP and FM 4-64-labeled hair cells in 5 dpf wild type and *lhfpl5b^vo35^* mutant *Tg(myo6b:GFP)vo68Tg* larvae. These images are from larvae that make up part of the data set quantified in Figure 4K. Scale bars = 5 µm.

**Supplemental Figure 5.**
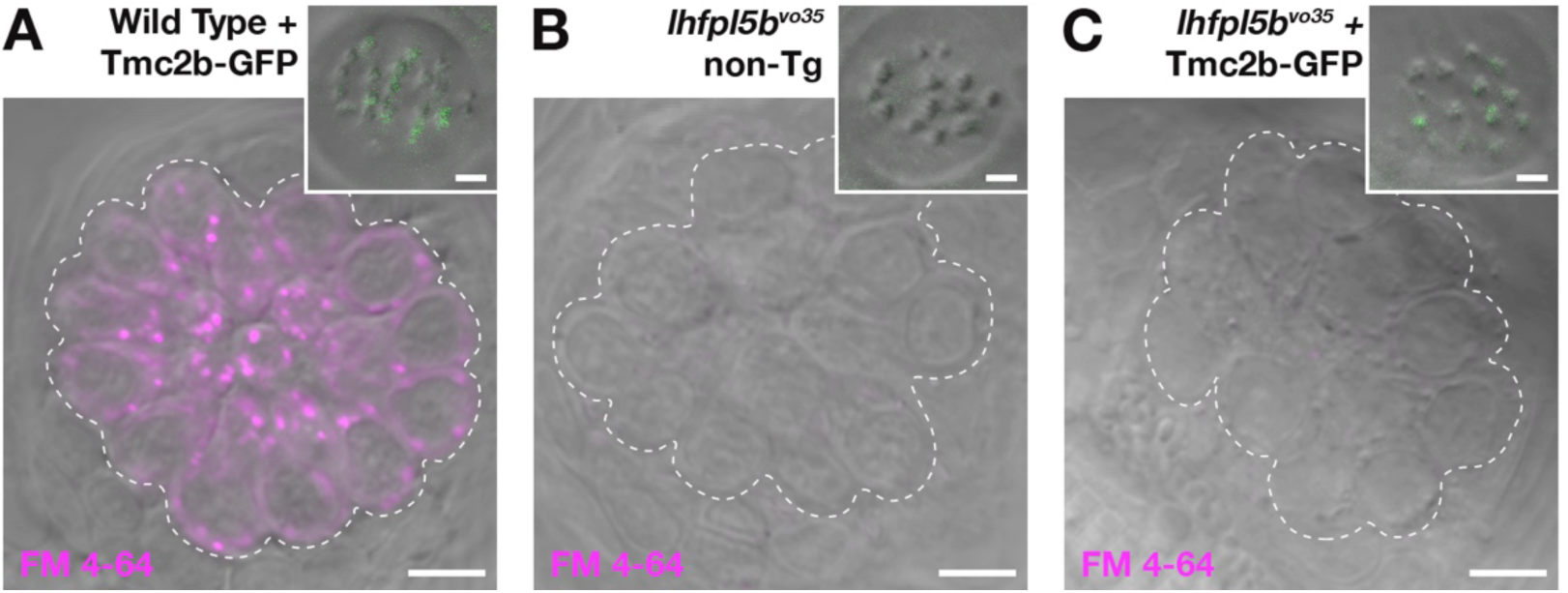
Bundle-localized Tmc2b-GFP does not restore basal MET channel activity to neuromast hair cells of *lhfpl5b^vo35^* mutants. (**A-C**) Representative images of neuromasts labeled with FM 4-64 from wild type Tmc2b-GFP *vo28Tg* (**A**), non-transgenic *lhfpl5b^vo35^* (**B**), and *vo28Tg*; *lhfpl5b^vo35^* (**C**) larvae at 7 dpf. Dashed lines outline the cluster of hair cells in each neuromast. Insets show the neuromast hair bundles from the same neuromast in the main panel. Scale bars = 5 µm in A-C; 2 µm for the bundle insets.

**Supplemental Figure 6.**
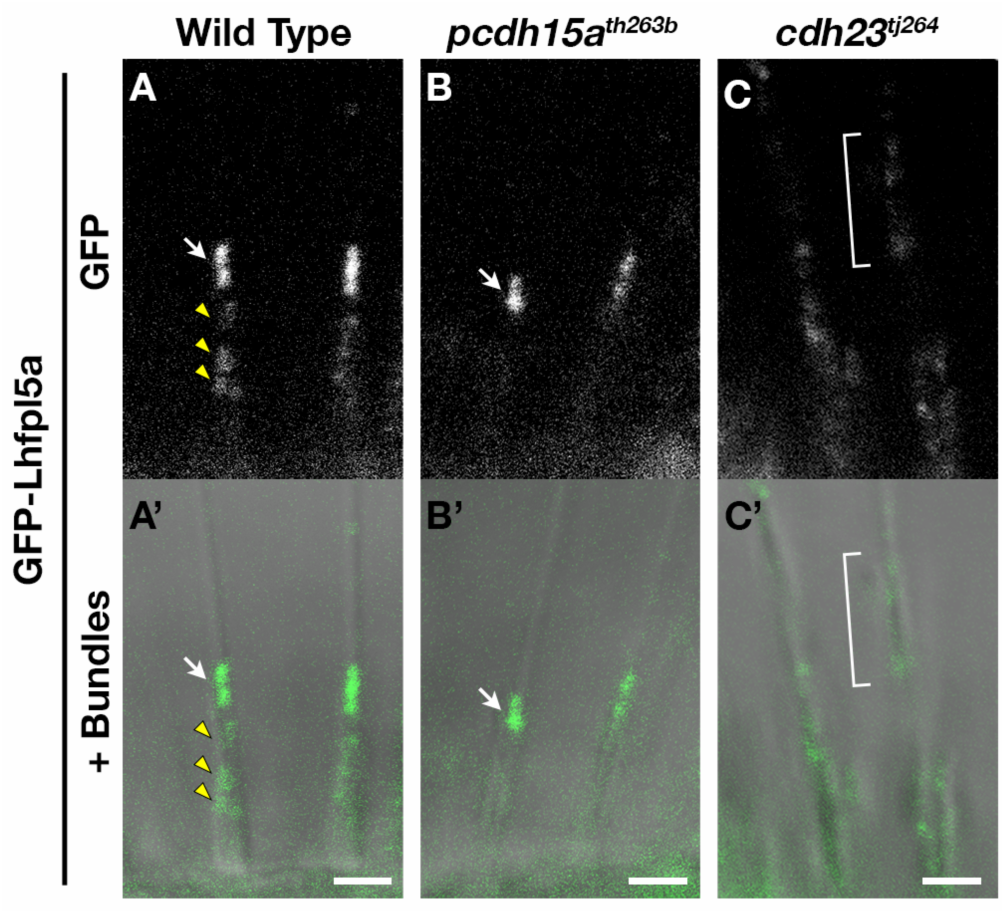
Mislocalization of GFP-Lhfpl5a in different mutant alleles of *pcdh15a* and *cdh23* as shown in Figure 6. Representative images of GFP-Lhfpl5a (*vo23Tg*) in the lateral cristae hair bundles of wild type (**A, A’**) and *pcdh15a^th263b^* (**B, B’**), and *cdh23^tj264^* (**C, C’**) mutants. The GFP-only channel is shown in panels A - C and overlaid with a light image of the bundles in A’ - C’. White arrows indicate GFP signal in the presumptive kinocilial linkages, yellow arrow heads indicate GFP signal in the stereocilia, and brackets indicate GFP signal in the kinocilium. Scale bars = 2 µm in A-C’.

